# Sex-specific transgenerational plasticity I: Maternal and paternal effects on sons and daughters

**DOI:** 10.1101/763862

**Authors:** Jennifer K Hellmann, Syed Abbas Bukhari, Jack Deno, Alison M Bell

## Abstract

1. Transgenerational plasticity (TGP) or parental effects – when parental environments alter the phenotype of future generations – can influence how organisms cope with environmental change. An intriguing, underexplored possibility is that sex –of both the parent and the offspring – plays an important role in driving the evolution of transgenerational plasticity in both adaptive and nonadaptive ways.
2. Here, we evaluate the potential for sex-specific parental effects in a freshwater population of threespined sticklebacks (*Gasterosteus aculeatus*) by independently and jointly manipulating maternal and paternal experiences and separately evaluating their phenotypic effects in sons versus daughters. We tested the adaptive hypothesis that daughters are more responsive to cues from their mother, while sons are more responsive to cues from their father.
3. We exposed mothers, fathers, or both parents to visual cues of predation risk and measured offspring antipredator traits and brain gene expression.
4. Predator-exposed fathers produced sons that were more risk-prone, while predator-exposed mothers produced more anxious sons and daughters. Further, maternal and paternal effects on offspring survival were nonadditive: offspring with a predator-exposed father, but not two predator-exposed parents, had lower survival against live predators. There were also strong sex-specific effects on brain gene expression: exposing mothers versus fathers to predation risk activated different transcriptional profiles in their offspring, and sons and daughters strongly differed in the ways in which their brain gene expression profiles were influenced by parental experience.
5. We found little evidence to support the hypothesis that offspring prioritize their same-sex parent’s experience. Parental effects varied with both the sex of the parent and the offspring in complicated and nonadditive ways. Failing to account for these sex-specific patterns (e.g., by pooling sons and daughters) would have underestimated the magnitude of parental effects. Altogether, these results draw attention to the potential for sex to influence patterns of TGP and raise new questions about the interface between transgenerational plasticity and sex-specific selective pressures, sexual conflict, and sexual selection.

## Introduction

Sex differences in life-histories (e.g. reproductive lifespan, mortality rate) or reproductive tactics can favor different optimal phenotypes in males and females (Andersson 1994). Although a shared genetic basis can constrain phenotypic differences between the sexes (Lande 1980; Reeve & Fairbairn 2001), epigenetic changes can overcome this constraint and allow males and females to respond differently to the same environmental condition (within-generational plasticity). Potentially adaptive sex-specific patterns of within-generational plasticity have been documented in diverse taxa (Stillwell *et al*. 2010; Ceballos & Valenzuela 2011; Xu *et al*. 2014; Meuthen *et al*. 2018); for example, in cichlids, predation risk experienced early in life influenced phenotypic development in males, but not females, possibly because males are more vulnerable to predation (Meuthen *et al*. 2018).

While less explored, there also is evidence for sex-specific ***transgenerational*** plasticity (TGP; also referred to as intergenerational plasticity or environmental parental effects); specifically, the sex of the parent and/or the offspring can alter the ways in which environments encountered by recent ancestors affect future generations. Empirical studies and theory to date have primarily focused on maternal effects; however, the biological reality is that the environment experienced by both mothers and fathers can affect future generations. For example, there is growing evidence for *paternal* effects on ecologically-important traits, which can be transmitted via paternal care as well as epigenetic changes to sperm (reviewed in (Crean & Bonduriansky 2014; Immler 2018)). Because males and females often experience different environments once they reach reproductive age and have different means of transmitting environmental cues to offspring (e.g. eggs versus sperm), the information transmitted by fathers may not match the information encoded by mothers. Indeed, there is mounting empirical evidence that maternal versus paternal exposure to the same environmental condition can have different effects on offspring (Bonduriansky & Head 2007; Bonduriansky, Runagall-McNaull & Crean 2016; Gilad & Scharf 2019). For example, previous studies have found both overlapping and distinct gene expressions of maternal versus paternal experiences (Beemelmanns & Roth 2016), which suggests that mothers and fathers can activate similar and different developmental programs in their offspring depending on their experience. Further, the influence of maternal environments might depend on paternal environments (or vice versa) (Mashoodh *et al*. 2012; Mashoodh *et al*. 2018; Zirbel & Alto 2018; Gilad & Scharf 2019); for example, a recent study by Lehto and Tinghitella (2020) found that stickleback females preferred duller males when either their mother or father had encountered predation risk, but preferred brighter males when both parents had experienced predation risk. Consequently, careful experimental studies that independently and jointly manipulate maternal and paternal effects are needed to understand the proximate and ultimate causes of similarities and differences between maternal and paternal effects.

Parental effects also often depend on the sex of the offspring. For example, parental environments can have opposing effects on the same trait in sons compared to daughters (Mueller & Bale 2007; Short *et al*. 2016; Braithwaite *et al*. 2017) or can influence different traits in sons versus daughters (Schulz *et al*. 2011; Metzger & Schulte 2016). For example, previous studies have found divergent gene expression in sons and daughters in response to maternal experiences (Metzger & Schulte 2016; Constantinof *et al*. 2019), which suggests that maternal experience activates different developmental programs in sons and daughters. Because the vast majority of studies that have compared parental effects on sons and daughters have focused on maternal effects, rather than both maternal and paternal effects (but see (Priest, Mackowiak & Promislow 2002; He *et al*. 2016; Emborski & Mikheyev 2019; Wylde *et al*. 2019)), it is often unclear if sex-specific offspring effects are driven by 1) differences in the magnitude of sons’ versus daughters’ responses to parental environments (e.g., daughters are generally more responsive to parental stress, whether mediated by the mother or the father) or 2) how offspring attend to experiences of their same-sex versus opposite-sex parent. Offspring may attend to experiences of their same-sex parent (Bouwhuis, Vedder & Becker 2015; Schroeder *et al*. 2015), perhaps because sex differences in life history strategies or dispersal result in daughters being more likely to encounter the environments experienced by their mothers, and sons being more likely to encounter their fathers’ environmental pressures. In order to evaluate the possibility that offspring selectively prioritize experiences of one parent over the other, it is necessary to compare maternal and paternal effects on both sons and daughters.

Sex-specific TGP might have important adaptive implications if it can resolve evolutionary conflicts that occur when selection favors different phenotypes in males and females (Bonduriansky & Day 2008). Mothers and fathers may selectively alter the phenotypes of their sons and daughters in response to the environment with a greater degree of precision than genetic inheritance and in ways that may match the distinct life-history strategies of males and females. Alternatively, sexual conflict could result in complex nonadaptive sex-dependent patterns, especially when sexual selection is strong (Burke, Nakagawa & Bonduriansky 2019). Here, we evaluate the potential for sex-specific TGP in threespined sticklebacks (*Gasterosteus aculeatus*). Male and female sticklebacks are sexually dimorphic in several respects, including in habitat use (Reimchen 1980), diet (Reimchen & Nosil 2001), parasite load (Reimchen & Nosil 2001), and morphology (Reimchen, Steeves & Bergstrom 2016), with these differences emerging during early adulthood (Reimchen 1980; Reimchen & Nosil 2001). Sexual selection favors a variety of male-specific reproductive traits that can increase males’ vulnerability to predation risk (Candolin 1998; Johnson & Candolin 2017): male sticklebacks develop bright nuptial coloration, engage in conspicuous territory defense and courtship behavior, and are the sole providers of paternal care that is necessary for offspring survival (Bell & Foster 1994). These sex differences in behavior and life history often expose males and females to different predation regimes (Reimchen & Nosil 2004), likely altering the environment experienced by mothers versus fathers and the optimal phenotype for daughters versus sons in response to predation risk.

We test the adaptive hypothesis that sex differences in life history strategies cause offspring to attend to cues from their same-sex parent: we predicted that daughters would attend to maternal cues and sons to paternal cues (i.e. a detectable interaction between maternal/paternal treatment, and offspring sex). To test this hypothesis, we exposed adult male and female sticklebacks to simulated predation risk prior to fertilization and used a fully factorial design to generate offspring of control (unexposed) parents, offspring of predator-exposed mothers, offspring of predator-exposed fathers, and offspring of predator-exposed mothers and fathers (Figure 1). Because predation risk varies in both space and time, it is likely that there is a mix of reproductively mature males and females who either have or have not recently experienced predation risk within many natural populations. We reared sons and daughters under ‘control’ conditions (i.e. in the absence of predation risk) and evaluated traits relevant to predator defense. We then evaluated offspring brain gene expression patterns to assess whether maternal experience with predation risk activates different developmental programs in offspring than paternal experiences, and whether the experience of one parent (e.g. fathers) activates a particular developmental program in one offspring sex but not the other (e.g. in sons but not daughters).

**Figure 1:**
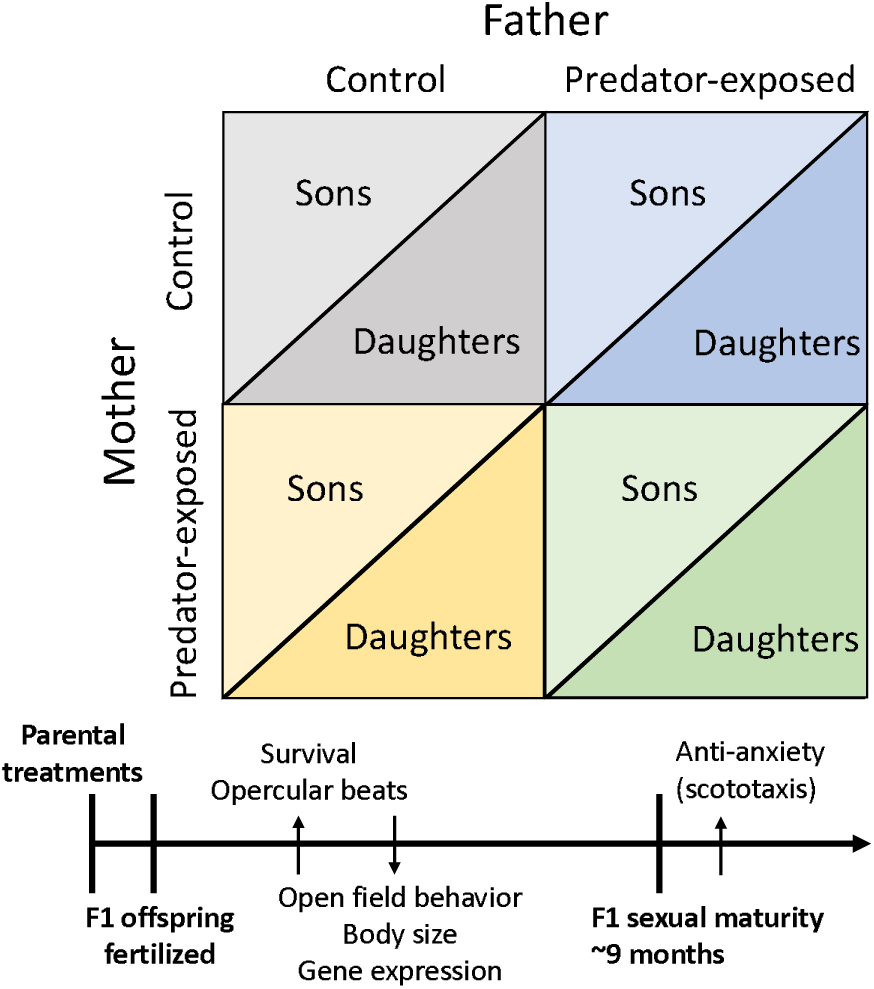
Experimental diagram of parental treatments with experimental timeline. Both male and female F0s were either left undisturbed or exposed to predation risk prior to fertilization and then crossed in a full-factorial, split-clutch design to generate F1 sons and daughters. On the timeline, bolded words represent key events while non-bolded words represent experimental measurements. Distinct sets of F1 offspring were used to measure 1) survival and opercular beats, 2) open field behavior and body size, 3) gene expression, and 4) anti-anxiety behavior (scototaxis).

## Methods

### Housing conditions

Adult, sexually-mature, freshwater threespined sticklebacks were collected from Putah Creek (CA, USA) in June-September 2016. This population has prickly sculpin (*Cottus asper*), which preys primarily on stickleback eggs, fry, and juveniles. To simulate natural conditions on the breeding grounds, where males defend nesting territories while females shoal, we used different procedures for exposing mothers and fathers to predation risk. Females were housed in six groups of n=10 fish per tank to mimic shoaling conditions in the wild. To simulate predation risk, we used a clay model sculpin (21cm long) to chase females for 90 seconds each day; unexposed treatment tanks were left undisturbed (similar to Dellinger *et al*. (2018)). Gravid females were removed from tanks and stripped of their eggs for in-vitro fertilization; this occurred the day after the last exposure for predator-exposed females. Control females were in the tank for the equivalent period of time and removed at the same time of day as the predator-exposed females to generate split clutches. Mothers were chased between 16-44 days, as mothers took a variable amount of time to become gravid (see Supplementary Material for additional analyses about the length of maternal exposure).

Males were housed singly to build nests. Once their nest was completed, predator-exposed males were chased by a model sculpin for 30 sec every other day for 11 days; control males were left undisturbed. A separate experiment confirmed that the results reported below were not produced when fathers were chased with a net (Chen *et al*. 2020), suggesting that changes in offspring traits are specific to predation risk and not a byproduct of, for instance, differences in activity levels due to chasing. The day after the last exposure, males were euthanized to obtain sperm for *in-vitro* fertilization. While female sticklebacks produce clutches of eggs repeatedly throughout the breeding season, stickleback males produce sperm in the beginning of the breeding season (Borg 1982); thus, paternal experiences mediated via sperm in this experiment are likely due to modifications to mature sperm. We used a short stressor to avoid reducing sperm quality or fertilization rates by exposing males to a stressor while developing sperm and to avoid potential habituation to predation risk (Dellinger *et al*. 2018). By beginning the treatment when males were transferred to a nesting arena, we sought to mimic the change in predator regime that males may encounter as they move into a different habitat to nest.

While females were exposed to risk in groups, males were exposed to risk singly. Because this regimen resulted in higher per capita risk for males than for females, males were exposed for a shorter period of time than females. Although mothers and fathers were exposed to risk differently, previous studies in this population suggest that exposure to predation risk for individuals who are isolated versus in groups have largely similar consequences: another experiment comparing post-fertilization paternal cues of predation risk with early life cues of predation risk found largely overlapping effects of paternal and personal experience with predation risk, despite the fact that fathers were exposed singly (with only one exposure) and offspring were exposed in family groups repeatedly for a week (Stein, Bukhari & Bell 2018).

F1 offspring were generated using a split clutch design, resulting in: 1) offspring of unexposed fathers and mothers (n=11 half-clutches), 2) offspring of exposed fathers and unexposed mothers (n=11 half-clutches), 3) offspring of unexposed fathers and exposed mothers (n=10 half-clutches), and 4) offspring of exposed fathers and mothers (n=10 half-clutches). By artificially fertilizing the eggs and incubating the embryos using an air bubbler, we controlled for possible pre-fertilization effects mediated by interactions between mothers and fathers (Mashoodh *et al*. 2012; McGhee *et al*. 2015), post-fertilization effects mediated by paternal care (Stein & Bell 2014), and the possibility that predator-exposed parents might be less likely to successfully mate or parent offspring. Separate groups of offspring were used for each assay described below (Figure 1; see Supplementary Material for more detailed methods).

### Measuring survival under predation risk and ventilation rate

At 3-5 months of age (mean: 20.6mm standard length ± 2.1 mm s.d), groups of n=4 offspring (one from each parental treatment) were exposed to a live sculpin predator. One day prior to the predation assay, fish were weighed, measured, marked, and individually transferred to a 250ml opaque glass beaker containing 100mL of water, with opaque sides to isolate the fish. We measured opercular beats 30 seconds after transferring to the beaker as a proxy for acute stress (Bell, Henderson & Huntingford 2010) and 30 minutes after transferring to understand response to prolonged stress (n=100 fish per parental treatment group). At the end of thirty minutes, all four fish were moved to the same holding tank until the predation trial the following day. For the predation trial, sticklebacks were simultaneously transferred into the sculpin’s tank (n=4 different sculpin, each used once per day); the trial ended two minutes after the first fish was captured by the sculpin. 14/100 trials did not result in any successful captures and were excluded from further analysis of survival data. We euthanized the survivors of the predation assays and used a section of muscle tissue to sex a large portion of the survivors per the methods of Peichel *et al*. (2004). We used generalized linear mixed models (GLMM) with a binomial distribution (R package lme4 (Bates *et al*. 2015)) to analyze differences in survival during the predation assay. We included fixed effects of maternal treatment, paternal treatment, and standard length, with random effects of maternal identity, paternal identity, sculpin identity, test group, and experimental day, to account for potential improvement in sculpin performance over time. Because we found evidence of heteroskedasticity in our opercular beat data, we used MCMC generalized linear mixed models (R package MCMCglmm (Hadfield 2010)) with a weak prior on the variance (V=1, nu=0.002) to analyze stress-induced respiration (breaths/minute). We ran models for 200,000 iterations, with a burn-in of 3000 iterations, thin = 3, and Gaussian distributions (and used these same parameters all MCMC models below). We included fixed effects of maternal treatment, paternal treatment, time period (30s or 30min), and standard length, as well as random effects of individual identity nested within both maternal and paternal identity. We removed one extremely low outlier in the opercular beat dataset. For all models, here and below, we tested for potential interactions between maternal treatment, paternal treatment, and offspring sex during our model selection process, and removed all not statistically significant interactions from final models.

### Measuring risk taking behavior

When offspring were 4.5 months, we measured behavior in an open field before and after a simulated predator attack (as in Bensky *et al*. (2017)). Individuals were placed in an opaque refuge in the center of a circular arena (150cm diameter) divided into eight peripheral sections with a circular section in the middle. After a three minute acclimation period, we removed the plug from the refuge, allowed fish to emerge, and then measured the number of different (exploration) and total (activity) sections visited for three minutes after emergence. We then simulated a sculpin predator attack; this attack elicited freezing behaviour from the fish and we measured the latency to resume movement after the simulated attack. Once the individual resumed movement, we again measured the number of different and total sections visited for three minutes. We weighed and measured the fish, euthanized it via decapitation, and preserved the body in ethanol for identification of sex (Peichel *et al*. 2004). We assayed n=118 fish: n=12 females and n=18 males with control parents, n=15 females and n=16 males with predator-exposed fathers, n=13 females and n=14 males with predator-exposed mothers, and n=11 females and n=19 males with two predator-exposed parents.

We used principal components analysis (R package factoextra (Kassambara & Mundt 2017)) to combine exploration and activity (Spearman rank correlation: ρ=0.92, p<0.001), using data from both before and after the simulated predator attack (two data points per individual). We extracted an eigenvalue of 1.77 that captured 88.4% of the variance in these two behaviors; positive values indicate more active and exploratory individuals. To understand how parental exposure to predation risk altered offspring activity/exploration, length, and body mass, we used MCMC GLMMs with a Gaussian distribution; we used MCMC GLMMs with a Poisson distribution to analyze offspring freezing behavior. All models include fixed effects of maternal treatment, paternal treatment, and individual sex. For activity/exploration, we included additional fixed effects of standard length, and observation period (before or after the simulated predator attack) and random effects of ID nested within mother and father identity (separately) and observer identity. For freezing behavior we included additional fixed effects of standard length and random effects of mother identity, father identity, and observer identity. We added an additional fixed effect of age (days since hatching) to the model testing length and an additional fixed effect of length for the model testing mass, with random effects of maternal and paternal identity for both models. We removed one outlier for the length dataset and one different outlier for the mass dataset.

### Measuring anxiety/cautiousness

Scototaxis (light/dark preference) protocols have been developed to test anti-anxiety/cautious behavior in fish (Maximino *et al*. 2010). When offspring were between 9-13 months old, offspring were gently caught with a cup from their home tank and placed in a clear cylinder (10.5cm diameter) in the center of a half-black, half-white tank (51L x 28W x 19H cm, coated on the inside and outside with matte contact paper). After a 5-minute acclimation period, we lifted the cylinder, and fish explored the tank for 15 minutes, during which we measured the latency to enter the white section, total time in the white section, and the number of times the fish moved between the black/white sections. After the 15-minute testing period, we removed the fish from the tank, recorded mass and standard length, euthanized the fish in an overdose of MS-222, and confirmed sex via dissection and examination of the gonads. The orientation of the tank was rotated between trials, and water was completely changed between each trial.

On average, fish spent less time in the white portion of the tank than the black portion (mean ± s.e.: 208.7 ± 18.8 sec out of a 900 sec trial). For individuals that never entered the white side of the tank (n=30 of 162 individuals), we recorded latency to enter as 900 seconds. Because all three variables were highly correlated (Spearman rank correlation; latency vs. time in white: ρ=-0.59, p<0.001; latency vs. movement between sides: ρ=-0.61, p<0.001; time in white vs. movement between sides: ρ=0.78, p<0.001), we used principal components analysis (R package factoextra) to combine these behaviors into one principal component (eigenvalue 2.10, captured 70.1% of the variance in behaviors): positive values were a measure of increased anti-anxiety/more cautious behavior with a higher latency to enter the white section, less total time in the white section, and fewer instances of switching between the black and white sections. We then used this principal component as the dependent variable in a MCMC GLMM with a Gaussian distribution, with fixed effects of maternal treatment, paternal treatment, sex, standard length, and day, to control for any potential season effects. We also included random effects of maternal and paternal identity, as well as observer identity. We assayed n=162 fish: n=23 females and n=15 males with control parents, n=22 females and n=17 males with predator-exposed fathers, n=23 females and n=21 males with predator-exposed mothers, and n=24 females and n=17 males with two predator-exposed parents.

### Measuring brain gene expression

We dissected whole brains from 4.5 month juvenile offspring We sampled one male and one female offspring per family from 5 families per treatment group, which totaled to n=10 (5 males, 5 females) offspring per treatment group (n = 40 total). We extracted RNA using Macherey-Nagel NucleoSpin 96 kits and sent n=39 samples to the Genomic Sequencing and Analysis Facility at UT Austin for TagSeq library preparation and sequencing (one sample was of poor quality). To estimate differential expression, pairwise comparisons between the experimental conditions (offspring with only a predator-exposed mother, offspring with only a predator-exposed father, offspring of predator-exposed mothers and fathers) relative to the control condition (offspring of unexposed parents) within each sex were made using edgeR (Robinson, McCarthy & Smyth 2010). To call differential expression, we used a ‘glm’ approach and adjusted actual p-values via empirical FDR, where a null distribution of p-values was determined by permuting sample labels for 500 times for each tested contrast and a false discovery rate was estimated (Storey & Tibshirani 2003).

In a separate analysis, we used WGCNA (R package WGCNA (Langfelder & Horvath 2008) to cluster genes into co-expressed gene modules (Zhang & Horvath 2005) and to reduce the dimensionality of the transcriptomic dataset, which allowed us to explore the potential for interactive effects of maternal treatment, paternal treatment and offspring sex on modules of genes with correlated expression patterns. To find modules associated with treatment effects, we fitted a linear model (Kuznetsova, Brockhoff & Christensen 2017) which blocked for clutch ID as random factor, along with main and interactive effects of sex, paternal treatment, and maternal treatment on module eigengenes. Eigengenes which were significantly associated (p < 0.05) with either the main or interactive effects of sex, paternal treatment, and maternal treatment were retained. See supplementary material for additional methods.

### Animal welfare note

All methods were approved by Institutional Animal Care and Use Committee of University of Illinois Urbana-Champaign (protocol ID 15077), including the use of live predators.

## Results

### Sons, but not daughters, of predator-exposed fathers were more active under risk

In the open field assay, offspring were less active/exploratory after the simulated predator attack compared to before (principal component analysis: higher values indicate more active and explorative individuals; Table 1), confirming that offspring behaviorally responded to the predator attack. There was a statistically significant interaction between paternal treatment and offspring sex on offspring activity/exploration (Table 1; Figure 2A). Specifically, sons of predator-exposed fathers were more active/exploratory compared to sons of control fathers (MCMC GLMM, 95% CI in brackets here and below [-1.30, -0.20], p=0.01), but there was not a detectable effect of paternal treatment on female offspring ([-0.40, 0.81], p=0.49). In other words, paternal effects on activity/exploratory behavior were stronger in sons, which is consistent with our hypothesis.

**Table 1:** Results of general linear mixed models (MCMCglmm) testing predictors of exploration/activity (higher values indicate more active and exploratory individuals) and freezing behavior in the open field assay, as well as anxiety-like behavior in the scototaxis assay. We tested for potential interactions between maternal treatment, paternal treatment, and offspring sex; we removed not statistically significant interaction terms.

|  | **Activity and exploration** |  |  |
| --- | --- | --- | --- |
|  | *Mean* | *95% CI (L, U)* | *P* |
| Observation period | -0.97 | -1.24, -0.70 | **<0.001** |
| Maternal treatment | 0.14 | -0.27, 0.54 | 0.48 |
| Paternal treatment | -0.20 | -0.81, 0.42 | 0.52 |
| Offspring sex | -0.25 | -0.70, 0.22 | 0.29 |
| Standard length | -0.12 | -0.18, -0.05 | **<0.001** |
| Paternal treatment * sex | 0.91 | 0.25, 1.54 | **0.005** |
|  | **Freezing behavior** |  |  |
|  | *Mean* | *95% CI (L, U)* | *P* |
| Maternal treatment | 0.31 | -0.15, 0.80 | 0.19 |
| Paternal treatment | -0.17 | -0.65, 0.31 | 0.47 |
| Offspring sex | -0.36 | -0.78, 0.07 | 0.09 |
| Standard length | 0.06 | -0.02, 0.15 | 0.14 |
|  | **Anxiety-like behavior** |  |  |
|  | *Mean* | *95% CI (L, U)* | *P* |
| Maternal treatment | 0.58 | 0.07, 1.11 | **0.03** |
| Paternal treatment | -0.19 | -0.70, 0.30 | 0.44 |
| Offspring sex | -0.72 | -1.27, -0.17 | **0.01** |
| Standard length | -0.05 | -0.10, -0.005 | **0.03** |
| Experimental day | 0.004 | -0.004, 0.01 | 0.33 |

**Figure 2:**
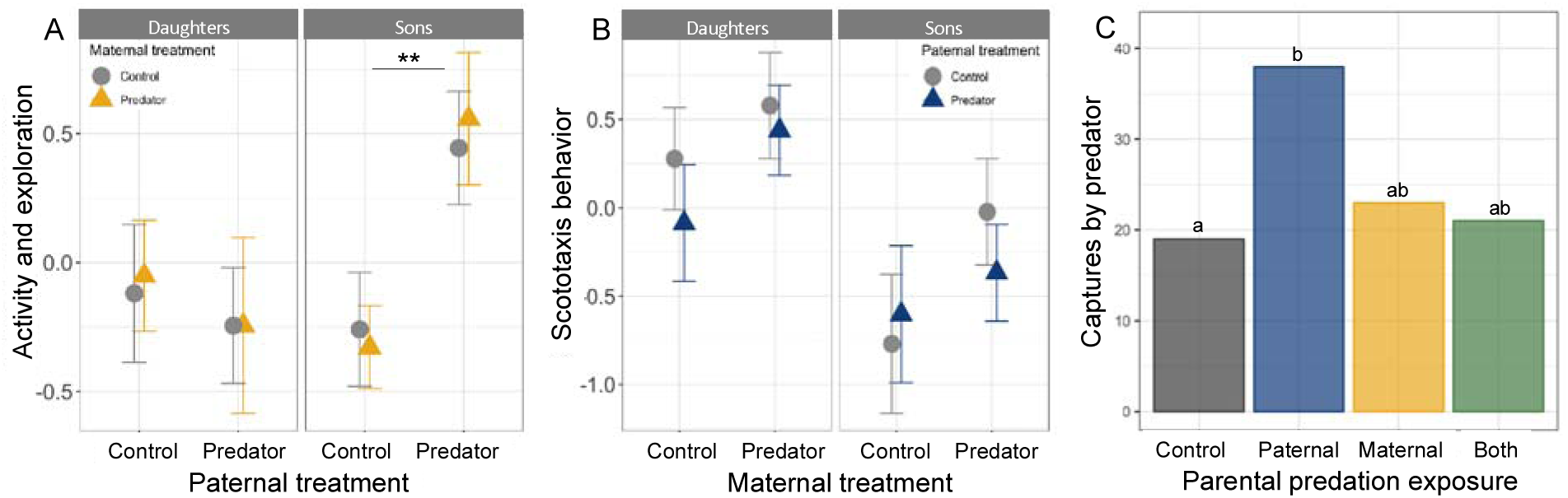
The effects of maternal and paternal treatment on offspring in an open field assay, scototaxis assay, and survival in the face of a live predator. A) Male offspring (right) of predator-exposed fathers were significantly more exploratory and active (PCA: higher values indicate more active and exploratory individuals; mean ± s.e.) compared to male offspring of control fathers; paternal treatment did not affect the exploratory behavior/activity of female offspring (left). The effect of paternal treatment did not depend on maternal treatment (control: grey; predator-exposed: yellow). N= 118 offspring. Stars indicate statistically significant differences across treatment groups. B) Offspring of predator-exposed mothers were more cautious (PCA: high values indicate longer latency to enter the white area and spent less time in the white area; mean ± s.e.) compared to offspring of control mothers. Further, female offspring (left) were more cautious than male offspring (right). The effect of maternal treatment did not depend on paternal treatment (control: grey; predator-exposed: blue). N= 162 offspring. C) In live predation trials, juvenile offspring of predator-exposed fathers, but not two predator-exposed parents, were more likely to be captured and consumed by the sculpin predator relative to offspring of control fathers. Shown are the total offspring captured across n= 86 trials; letters indicate significant differences among treatment groups, determined by Tukey’s HSD with parental treatment as a 4-level variable.

We did not detect any maternal or paternal effects on freezing behavior (Table 1) or standard length or body mass (see Supplementary Table 1). However, while we found no overall differences between offspring of control and predator-exposed mothers in length, among offspring that had a predator-exposed mother, longer maternal exposure to the predator resulted in larger offspring at 4.5 months (Supplementary Results). The length of exposure did not significantly alter any other measured offspring traits (see supplementary material), although there may have been an effect on other traits that were not measured in this study. Standard length, mass, and freezing behavior (Table 1, Supplementary Table 1) also did not vary between male and female offspring, although larger fish were less active/exploratory (Table 1).

### Both sons and daughters of predator-exposed mothers, but not fathers, were more cautious

Offspring of predator-exposed mothers were more cautious (principal component analysis: took longer to enter the white area, spent less time in the white area, and switched less between black and white areas) compared to offspring of control mothers (Table 1; Figure 2B). However, we did not detect an effect of paternal treatment on offspring scototaxis behavior (Table 1). Both female and smaller offspring showed more cautious behavior, and we found no evidence of seasonal effects (Table 1). Consequently, rather than offspring attending to the experiences of their same-sex parent in the scototaxis assay, we observed that both sons and daughters responded to maternal experiences.

### Offspring of predator-exposed fathers were more vulnerable to predation, but not if their mother was also exposed

There was a statistically significant interaction between maternal and paternal treatment on offspring survival in live predation assays (generalized linear mixed effect model: Z_334_ = - 1.98, 0.048). Specifically, offspring of predator-exposed fathers were more frequently captured by the predator compared to offspring of control parents (Tukey’s HSD with parental treatment as a 4-factor variable: Z=2.71, p=0.03), but this was not true for offspring of predator-exposed mothers (Z=0.75, p=0.88) or both a predator-exposed mother and father (Z=-0.81, p=0.85; Figure 2C). These results suggest that there was a strong fitness cost of having a predator-exposed father, but mothers seemed to mitigate those costs, perhaps by making their offspring more cautious (see above). Survivors of the successful predation trials were heavily female biased (93/148; Chi-squared: χ^2^=9.76, p=0.002), suggesting that males are generally more vulnerable to predation risk. The sex-bias was not statistically different across treatment groups (χ^2^=3.03, p=0.39); this suggests that paternal exposure influenced sons and daughters equally, although we cannot conclude this definitively because we do not know the sex of the captured fish. We found no effect of size on how frequently the stickleback were captured by the predator (Z_334_ = 1.56, 0.12).

We did not detect any maternal or paternal effects on stress-induced respiration (see Supplementary Table 1). For the portion of offspring where sex was known, we did not detect any interaction between offspring sex and paternal ([-17.67, 14.83], p=0.89) or maternal treatment ([-23.89, 8.77], p=0.37), although males tended to have higher opercular beats than females (main effect of sex [-1.48, 25.92], p=0.08).

### Distinct maternal and paternal effects on offspring brain gene expression

To evaluate whether predation risk experienced by mothers versus fathers has different consequences for offspring development at the molecular level, we compared the baseline brain gene expression profile of offspring of unexposed parents (control) to offspring with a predator-exposed mother, a predator-exposed father, and two predator-exposed parents in male and female offspring (n=39 individuals). In terms of the number of genes, maternal and paternal effects on brain gene expression were approximately equivalent in magnitude, and the genes were largely nonoverlapping (Figure 3A,B): in sons, for example, 1028 genes were differentially expressed in response to maternal experience with risk, 904 genes were differentially expressed in response to paternal experience with risk while only 253 genes were shared between them (daughters show a similar pattern, Figure 3A). This suggests that, in contrast to our prediction, the transcriptomes of sons and daughters are not more responsive to the experiences of their same-sex parent. Interestingly, there was also a large number of genes that were unique to the “both” condition, i.e. between offspring of two predator-exposed parents versus the control; these differentially expressed genes could reflect the ways in which maternal and paternal effects interact at the molecular level.

**Figure 3:**
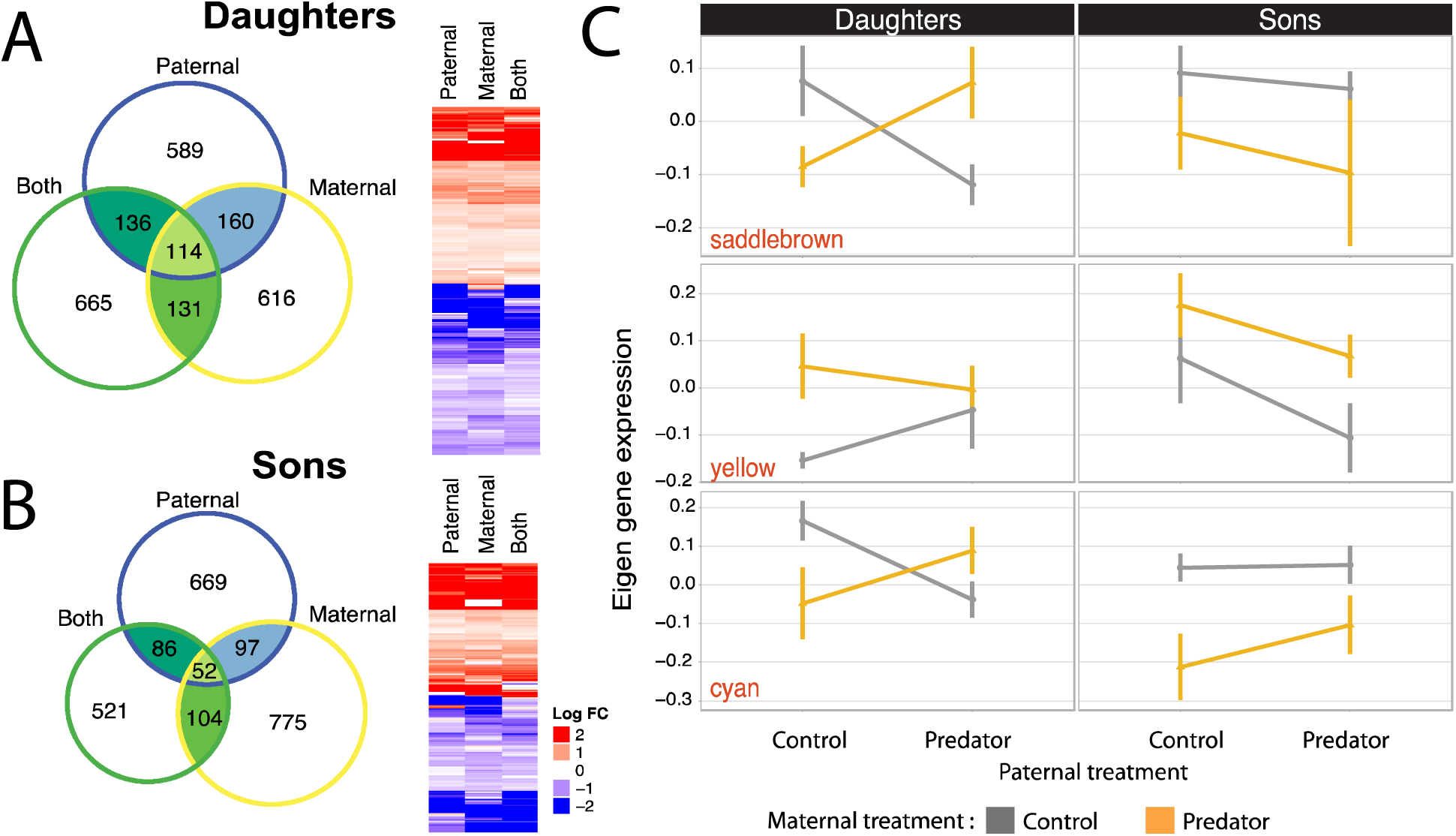
Differential gene and eigen-gene expression analysis. A-B) The three circles in the Venn diagram show the number of genes that were differentially expressed in the brain of offspring of unexposed parents relative to offspring of predator-exposed mothers (“maternal”), predator-exposed fathers (“paternal”), or two predator-exposed parents (“both”), with daughters in (A) and sons in (B). Note that relatively few genes overlap between the different pairwise comparisons. The heatmaps show the direction of gene regulation (blue: downregulated; red: upregulated) of the differentially expressed genes that are shared among the three pairwise comparisons, with daughters and sons shown separately. C) The expression profiles of the three eigen-gene modules which were affected by the three-way interaction among paternal treatment, maternal treatment and offspring sex (mean ± s.e.). N=39 offspring.

Of the differentially expressed genes that were shared between the pairwise comparisons, nearly all were concordantly regulated, for both sons and daughters (Figure 3A,B). This suggests that, despite the large-scale differences in brain gene expression between offspring of predator-exposed mothers and fathers, there is a core set of genes that is activated in offspring brains in response to either maternal or paternal exposure to predation risk.

### Maternal and paternal exposure to predation risk interacted with offspring sex to influence offspring brain gene expression

WGCNA identified 23 clusters (“modules”) of genes with coordinated expression patterns in the dataset. The expression of eight of the 23 modules was affected by at least one of the factors in the model: three modules were affected by maternal treatment, two were affected by the two-way interaction between maternal and paternal treatment, and three were affected by the three-way interaction between paternal treatment, maternal treatment and offspring sex (shown in Figure 3C). For example, the module “saddle brown” comprises 48 co-expressed genes (largely enriched for developmental processes) whose expression was influenced by the three-way interaction between maternal treatment, paternal treatment and offspring sex. Specifically, daughters of a predator-exposed mother or father showed lower expression of genes in this module compared to daughters of control parents or two predator-exposed parents (Figure 3C). For sons, on the other hand, the expression of genes in this module was more strongly affected by maternal treatment. A similar pattern was observed in the yellow and cyan modules. Overall these results demonstrate that at the molecular level, daughters and sons differ in the extent to which they respond to predation risk that had been experienced by their mother, father or by both parents. However, in contrast to our overall hypothesis, there was no evidence that sons and daughters primarily attend to experiences of their same-sex parent at the molecular level.

## Discussion

Here, we report the results of a comprehensive comparison of maternal and paternal effects on sons and daughters. We show that both the sex of the parent and the sex of the offspring influence the ways in which offspring phenotypes are altered by parental experiences. However, we found little evidence for the adaptive hypothesis that offspring would attend primarily to cues from their same-sex parent. Instead, our results illustrate the complexities of sex-specific parental effects. Maternal and paternal effects in response to the same environmental factor were largely distinct: predator-exposed mothers produced more cautious offspring (scototaxis), while predator-exposed fathers produced sons, but not daughters, that were more active under risk (open field assays). Further, the effects of paternal predation exposure were moderated by maternal predation exposure: offspring of predator-exposed fathers, but not two predator-exposed parents, had reduced survival against a live predator. Yehuda *et al*. (2014) found this same interactive pattern previously in humans due to parental PTSD and originally attributed it to potential changes in parenting behaviors. Our results demonstrate that these non-additive interactions can be mediated in the absence of behavioral interactions between parents and offspring. Instead, our brain gene expression profile results – that there are distinct neurogenomic changes in offspring with two predator-exposed parents compared to offspring of either a predator-exposed mother or father – are consistent with the hypothesis that non-additive patterns may arise because offspring phenotypes are influenced by the interaction between environmentally-induced epigenetic changes in eggs versus sperm.

Sex-specific patterns – where sons and daughters differ in their response to maternal and paternal exposure to predation risk – emerged in our study well before offspring were reproductively mature, during a period in their life when males and females are shoaling and occupying similar habitats (Bell & Foster 1994). In contrast to our prediction that offspring would adaptively attend to the cues of their same-sex parent, these sex-specific patterns of parental effects did not seem to emerge along a consistent male-female divide (e.g. sons attend to their father and daughters attend to their mother); instead, sons and daughters were both altered by paternal and maternal environments, but in different ways. This is consistent with Emborski and Mikheyev (2019), who found that male offspring were influenced by maternal, but not paternal, diet. These sex-specific effects may be adaptive for offspring, with differences originating in early development to allow offspring to develop phenotypes that are better matched to the different environments they will encounter later in life. For example, it is possible that increased activity under risk for sons, but not daughters, may be adaptive because high variance in male reproductive success favors males that adopt high risk, high reward behaviors to increase growth and access to resources under high predation pressure (Bell, Henderson & Huntingford 2010). Alternatively, these effects may not be adaptive, either resulting from differences in sons and daughters in their susceptibility to parental stress (Bale 2011; Glover & Hill 2012) and/or reflecting sexual conflict and sexually antagonistic selection (Burke, Nakagawa & Bonduriansky 2019). Indeed, Burke, Nakagawa and Bonduriansky (2019) suggest adaptive TGP may be unlikely to arise in systems with sex-specific selection because sex-specific ecologies can result in mothers and fathers experiencing different environments and therefore transmitting conflicting information to their offspring.

Our study shows that maternal and paternal predation exposure can have fitness consequences for offspring (i.e., via survival) in the lab; work is needed in a more natural context in the field to assess the fitness consequences of parental effects. For example, there are multiple steps required to avoid predation (Lima & Dill 1990; Guiden *et al*. 2019); while our data suggest that offspring with predator-exposed fathers are poor at evading predators once they come into contact with predators, parental experience with predation risk might alter the likelihood that offspring initially avoid coming into contact with predators. Further, offspring of predator-exposed fathers might face a trade-off between survival and reproduction, favoring high-risk, high-reward strategies that reduce survival in high predation environments, but increase reproductive success by ensuring that surviving individuals are in good breeding condition. Indeed, sticklebacks do seem to face a trade-off in surviving predation and gaining the body size necessary for successfully reproducing, with this trade-off being stronger in males compared to females (Bell *et al*. 2011).

Whether the fitness interests of mothers, fathers, and offspring align or conflict has important implications for the evolution of sex-specific TGP (Burke, Nakagawa & Bonduriansky 2019). When parents’ and offspring fitness interests in the face of predation risk are aligned, sex-specific plasticity may arise because mothers and fathers experience their environment in different ways and/or because the same parental environment favors different phenotypes in sons and daughters. However, sex-specific TGP may arise because mothers and fathers favor different optimal offspring phenotypes (Saldivar *et al*. 2017), and/or sons and daughters have different capacities to respond to or ignore information from fathers and mothers. If this is the case, TGP may evolve at the interface between sexual conflict and parent-offspring conflict, with paternal strategies, maternal strategies, and offspring counter-adaptations all ultimately dictating offspring phenotypes. This may result in the evolution of mechanisms that allow mothers to manipulate the ways in which fathers influence offspring (e.g. via cytoplasmic contributions (Crean & Bonduriansky 2014)) or fathers to manipulate the ways in which mothers influence offspring (e.g. via ejaculate composition (Garcia-Gonzalez & Dowling Damian 2015)).

Interactions between maternal effects, paternal effects, and offspring sex could be mediated via a variety of proximate mechanisms. Distinct maternal and paternal effects could reflect different proximate mechanisms that mediate the transmission of cues from mothers versus fathers to offspring (e.g., egg hormones or mRNAs versus sperm small RNAs) as well as the ways in which mothers and fathers were exposed to risk. Both distinct and interactive effects could also be mediated by epigenetic mechanisms such as parent-of-origin effects (Kong *et al*. 2009; Lawson, Cheverud & Wolf 2013) or interactions between maternal and paternal contributions (e.g. egg cytoplasm altering the effect of sperm small RNAs) during early development (Crean & Bonduriansky 2014; Garcia-Gonzalez & Dowling Damian 2015).

Differences between sons and daughters in how they respond to parental information could be mediated via trans-acting mechanisms (e.g., regulation of genes on non-sex chromosomes by genes located on the sex chromosome (Metzger & Schulte 2016)), sex-specific differences in epigenetic mechanisms, or genomic imprinting (Bonduriansky & Day 2008; Dunn & Bale 2011). Further, in bulls, Y-bearing and X-bearing spermatozoa have differentially expressed proteins, suggesting a mechanism by which fathers can transmit different information to sons versus daughters (Scott *et al*. 2018). Although mothers in many species can also transmit different information to sons and daughters (e.g., via placental function and gene expression (Bale 2011; Glover & Hill 2012)), it is unclear if mothers can transmit different information to sons and daughters in externally fertilizing species such as sticklebacks, in which mothers do not interact with their offspring post-fertilization. Future work exploring these proximate mechanisms could help explain the extent to which variation in parental effects is due to changes in the information encoded by parents or changes in offspring responsiveness to parental information.

Because parents can differentially allocate based on their partner’s phenotype or environmental conditions experienced by their partner (Mashoodh *et al*. 2012; McGhee *et al*. 2015; Mashoodh *et al*. 2018), in most systems it is difficult to isolate the effects of direct parental exposure to an environmental cue from environmental cues that parents indirectly detect from their mate (e.g. predator-naïve fathers provide less care to offspring of predator-exposed mothers) (Mashoodh *et al*. 2012; McGhee *et al*. 2015; Mashoodh *et al*. 2018). This makes it difficult to understand whether paternal effects can be mediated via sperm alone, or to determine the influence of paternal effects in isolation of maternal effects. In this experiment, we were able to completely isolate paternal effects mediated via sperm because there was no opportunity for parents to interact pre-fertilization or to influence offspring post-fertilization. Although our results suggest that distinct and interactive effects of maternal and paternal effects can be mediated via selective changes to information encoded in eggs and sperm alone, a fascinating direction for future work would be to consider how parental care and mate choice might ameliorate or magnify the sex-specific effects observed here.

## Conclusions

Transgenerational plasticity can allow environmental information to be delivered to offspring earlier and with potentially lower costs to offspring than developmental plasticity (Bell & Hellmann 2019). TGP can potentially be fine-tuned to the precise environment that both parents and offspring will encounter (Bonduriansky & Day 2008), perhaps including the different environments experienced by males and females because of sex differences in life history and reproductive tactics. We found that paternal cues mediated via sperm seem to be just as prominent in terms of their magnitude and prevalence as maternal cues mediated via eggs, although differences in our exposure regime (due to stickleback biology) may have influenced the relative strength of maternal versus paternal effects. Furthermore, we show that offspring phenotypes varied depending on whether predation risk had been experienced by their mother or their father, and a parent’s experience with predation risk produced different phenotypes in their sons compared to their daughters. However, these sex-specific patterns would have been masked if we had combined cues coming from mothers and fathers (i.e. compared offspring of two predator-exposed parents to a control) or failed to isolate effects emerging in sons versus daughters. Consequently, theoretical and empirical work seeking to understand the evolution of transgenerational plasticity would benefit from considering the conditions which influence *sex-specific* patterns of transgenerational plasticity in both adaptive and nonadaptive ways. Further, given broad interest in understanding the consequences of transgenerational plasticity for future generations and its potential to influence adaptive evolution, future work should consider how sex-specific effects in the first generation may alter the ways in which intergenerational effects persist for multiple generations in lineage-specific and/or sex-specific ways.

## Acknowledgements

Thank you to Eunice Chen, Erin Hsiao, Yangxue Ma, Liam Masse, and Christian Zielinksi for help with data collection and to Sarah Donelan and the Bell lab for comments on previous versions of this manuscript. This work was supported by the National Institutes of Health award number 2R01GM082937-06A1 to Alison Bell and National Institutes of Health NRSA fellowship F32GM121033 to Jennifer Hellmann. The authors have no conflicts of interest.

## Author contributions

JKH and AMB designed the study. JKH generated offspring, conducted survival assays, collected opercular beat data, dissected brains and extracted RNA, and oversaw open field assays and offspring sexing. JD conducted scototaxis assays. SAB conducted gene expression analyses. JKH wrote the first draft of the manuscript and JKH/AMB edited the manuscript.

## Data accessibility

All datasets (survival, respiration data, scototaxis, behavioral assays, lists of differentially expressed genes, read counts per sample, WGCNA) will be made publicly available on Dryad upon acceptance of this manuscript.

