## Supplementary Material for "Sex-specific transgenerational plasticity I: Maternal and paternal effects on sons and daughters"

**Additional methodological details**

***Housing and rearing conditions*.** The F0 generation was maintained on a summer photoperiod schedule (16 L : 8D) at 21° ± 1°C and fed ad libitum daily with a mix of frozen bloodworms (*Chironomus* spp.), brine shrimp (Artemia spp.) Mysis shrimp, and Cyclop-eeze. To simulate natural conditions on the breeding grounds, where males defend nesting territories while females shoal together, we used different procedures for exposing mothers and fathers to predation risk. Mothers were housed in six groups of n=10 fish per tank (37.9L, 53 × 33 x 24 cm high) to mimic shoaling conditions in the wild. Fathers were kept singly in 26.5L tanks (36L x 33W x 24H cm), visually isolated from the other males’ tanks with opaque partitions. Each tank contained two plastic plants, a sandbox, a clay pot, and algae to encourage nest building.

F1 offspring were generated via *in vitro* fertilization using a split clutch design (August-November 2016). Each female’s clutch was split and fertilized with sperm from both a predator-unexposed and predator exposed male, while each male’s sperm was split and used to fertilized eggs from a predator-unexposed and predator exposed female, ultimately resulting in 42 successful clutches of half-siblings across the four parental treatment groups. Predation risk did not alter fertilization rates or clutch survival: in addition to the successful clutches listed above, we had 3 failed clutches of control parents, 2 failed clutches of predator-exposed fathers, 3 failed clutches of predator-exposed mothers, and 3 failed clutches of predator-exposed fathers and mothers.

We incubated fertilized eggs in a cup with a mesh bottom placed above an air bubbler and fry were reared in 37.9 L tanks (53L x 33W x 24H cm). Offspring were fed newly hatched brine shrimp for 2 months before transitioning to the same mix of frozen food noted above. At 2.5 months, we split the five largest clutches in each of the four treatments. We gently caught individual fish in a clear bottomed cup and put 10 fish each into two 26.5L tanks. Fish in one of those 26.5L tanks were used for the behavioral assays and gene expression (see below). In the remaining clutches, 20 fish were caught and immediately returned to their tank. Larval offspring were switched from a summer photoperiod schedule (16 L : 8D) at 21° ± 1°C to a winter light schedule (8 L: 16 D) at 20° ± 1°C prior to the predation trials, open field assays, and brain collection, but resumed a summer light schedule prior to the scototaxis assays, which were conducted when offspring were reproductively mature. Separate groups of offspring were used for each assay described below.

***Measuring survival under predation risk and ventilation rate****.* One day prior to the predation assay, one fish from each of the four parental treatments was gently caught from their home tank, weighed, and measured; fish within a trial were size matched as much as possible (mean pairwise standard length (SL) difference among stickleback per trial: 2.06mm ± 0.90 mm s.d). We gave each fish one mark with blue, yellow, orange, or red elastomer dye (Northwest Marine Technologies) on the side of their body. To control for potential correlations between color and survival, the color was rotated among trials such that all treatments received each color for one-fourth of the trials (captures did not vary by color: Chi-squared test, χ^2^=3.50, p=0.32). After marking, each fish was transferred to a 250ml glass beaker to measure opercular beat rate. At the end of thirty minutes, all four fish were moved to the same 9.5L holding tank (32 x 21 cm and 19 cm high) until the predation trial the following day.

Sculpin used as predators in this experiment (n=4) were housed individually in 26.5L tanks (36L x 33W x 24H cm) with a bubbler, plants, and a clay pot for shelter. Sculpin were fed with only live prey (namely guppies) for several months prior to the beginning of the trials to habituate them to capturing live prey in a laboratory setting. Each sculpin was used for a maximum of one trial per day. One hour prior to the beginning of the trial, all bubblers and plants were removed from the sculpin’s tank and water was drained to the halfway point. Immediately prior to the trial, the sticklebacks were gently netted from their holding tank, placed in water in individual cups, and simultaneously transferred into the sculpin’s tank as far away from the sculpin as possible. The trial commenced as soon as all four fish were in the testing tank and ended two minutes after the first fish was captured by the sculpin. We left the stickleback in the tank for up to three hours and recorded the identity of the survivors. We used a section of muscle tissue to sex a large portion of the survivors (n=157 fish from 67 trials; tissue samples were not collected for the first n= 22 trials) with a male-specific genetic marker per the methods of Peichel *et al.* (2004).

***Measuring behavior under predation risk.*** When offspring were 4.5 months, we measured behavior in an open field before and after a simulated predator attack (as in Bensky *et al.* (2017)). Individuals were placed in an opaque refuge and given a three minute acclimation period before we removed the plug from the refuge, allowed fish to emerge, and then measured the number of different (exploration) and total (activity) sections visited for three minutes after emergence. Fish that did not emerge from the refuge after 10 minutes were gently released from the refuge; while offspring who emerged naturally were more active/exploratory than fish who were released (generalized linear model with binomial distribution (emerged or released), with activity/exploration as a fixed effect: Z_234_=-3.68, p<0.001), controlling for emergence time did not alter the significance of the results reported below.

After the 3min period, we simulated a sculpin predator attack by briefly moving the model predator sculpin in the direction of the offspring for 5 seconds (it was not present in the arena prior to the attack and was removed from the arena after the attack). This attack elicited freezing behaviour from the fish; once the individual resumed movement, we again measured the number of different and total sections visited for three minutes. If the fish remained frozen for greater than five minutes (n=20 fish), we ended the trial and considered activity and exploration after the simulated predation attack to be zero. Statistics were conducted on all the data, but results remain significant when these fish are omitted.

***Measuring brain gene expression****.* Offspring were captured from their home tank between 1100-1600hrs and immediately sacrificed. Brains were preserved in RNAlater, stored at 4°C overnight, and transferred to -80°C until RNA extraction. We extracted RNA using Macherey-Nagel NucleoSpin 96 kits, confirmed quality of samples via Bioanalyzer, and normalized the concentration of the samples prior to library preparation and sequencing.

##### *Additional statistical Analysis*.

*Stress-induced respiration.* To determine if offspring sex predicted opercular beat rate, we reran the model described in the main text on the portion of offspring where sex was known (the survivors of the predation assays, n=157 offspring); we included all fixed and random effects stated above, plus an additional fixed effect of offspring sex. To determine if opercular beat rate predicted survival in the predation assays, we used GLMMs with a binomial distribution with survival as the dependent variable, opercular beat rate (square-root transformed) as a fixed effect, and random effects of maternal identity, paternal identity, sculpin identity, experimental day, and test group. We ran separate models for opercular beat rate at 30 seconds and 30 minutes.

*TagSeq informatics.* FASTQC was used to assess the quality of the reads. Tag-seq produced an average of ~7 million reads per sample. We aligned reads to the *Gasterosteus aculeatus* reference genome (the repeat masked reference genome, Ensembl release 92), using STAR (2.5.3) (Dobin *et al.* 2013). We assigned reads to features according to the Ensembl release 92 gene annotation file ([http://ftp.ensembl.org/pub/release-92/gtf/gasterosteus_aculeatus/)](http://ftp.ensembl.org/pub/release-72/gtf/gasterosteus_aculeatus/)). We used HTSeq-Count to count reads mapped to gene features using stickleback genome annotation. We excluded multimapped reads or reads mapped to non-genic location from the analysis.

*Differential gene expression.* We excluded wo samples based on high variability on multidimensional scaling (MDS) plots. We included genes with at least 0.1 cpm in 5 samples. To estimate differential expression, we made pairwise comparisons between control and each treatment group (offspring with just a predator-exposed mother, with just a predator-exposed father, or two predator-exposed parents) within each sex using edgeR. Count data were TMM (trimmed mean of M-values) normalized and we used a ‘glm’ approach to call differential expression between treatment groups. We adjusted actual p-values via empirical FDR, where a null distribution of p-values was determined by permuting sample labels for 500 times for each tested contrast and a false discovery rate was estimated (Storey & Tibshirani 2003).

*Co-expression network analysis.* To build an unsigned weighted co-expression network, we excluded genes with non-zero variances. Further, we excluded genes that had zero counts in at least 80% of the samples. We voom transformed the input counts using voom functionality (R package limma) and then estimated pairwise gene-gene correlations using Pearson correlation. Based on scale free topology criterion, we removed spurious correlations by estimating appropriate soft threshold. We used a soft threshold of 10 with almost 90% R square to filter low correlations and to build adjacency (A) and topological overlap (TOM) matrices. Then using hierarchical clustering, we built a dendrogram of all genes based on TOM. This tree was cut using dynamic tree cut method as implemented in WGCNA (Zhang & Horvath 2005; Langfelder & Horvath 2008) with deepSplit = 4 and minimum cluster size of 30 genes. Modules were further merged based on their similarity and their eigengenes values were saved for downstream analysis. Finally, we retained genes with > 0.5 correlations with module eigengenes as modules members.

To find modules associated with treatment effects, we fitted a linear model which blocked for clutch ID as a random factor, along with main and interactive effects of sex, paternal treatment, and maternal treatment on module eigengenes using lmer test function in lmerTest package in R (Kuznetsova, Brockhoff & Christensen 2017). We used clutch ID for this analysis, rather than maternal and paternal identity separately, because we selected offspring from a subset of clutches and few of the clutch shared mothers or fathers. Eigengenes which were significantly associated (p < 0.05) with either the main or interactive effects of offspring sex, paternal treatment, and maternal treatment were retained.

**Results**

***Opercular beats***. We found no detectable difference between stress-induced respiration rates after initial confinement or after 30 minutes of confinement (Supplementary Table 1). Larger fish tended to have lower stress-induced respiration compared to smaller fish (Supplementary Table 1). Individuals with lower opercular beat rate at initial confinement (Z_335_= -1.92, p=0.05) and after 30 minutes of confinement (Z_335_= -1.75, p=0.08) tended to be more likely to be captured by the predator. Given that we do not have a measure of baseline respiration rate, we do not know if these differences in stress-induced respiration derive from variation in baseline respiration rates or in responsiveness to the confinement stress.

**Supplementary Table 1:** Results of general linear mixed models (MCMCglmm) testing predictors of standard length and mass at 4.5 months, as well as stress-induced respiration. We tested for potential interactions between maternal treatment, paternal treatment, and offspring sex; we removed not statistically significant interaction terms.

|  | **Standard length** | | |
| --- | --- | --- | --- |
|  | *Mean* | *95% CI (L, U)* | *P* |
| Maternal treatment | 0.36 | -1.14, 2.02 | 0.66 |
| Paternal treatment | -0.19 | -1.72, 1.36 | 0.76 |
| Offspring sex | -0.46 | -1.27, 0.33 | 0.25 |
| Days since hatched | 0.11 | 0.06, 0.17 | **<0.001** |
|  | **Mass** | | |
|  | *Mean* | *95% CI (L, U)* | *P* |
| Maternal treatment | -0.001 | -0.03, 0.02 | 0.91 |
| Paternal treatment | -0.0005 | -0.03, 0.03 | 0.96 |
| Offspring sex | 0.005 | -0.003, 0.01 | 0.20 |
| Standard length | 0.02 | 0.019, 0.022 | **<0.001** |
|  | **Stress-induced respiration** | | |
|  | *Mean* | *95% CI (L, U)* | *P* |
| Maternal treatment | 3.67 | -4.78, 12.45 | 0.37 |
| Paternal treatment | -6.63 | -17.09, 3.93 | 0.20 |
| Observation period | 0.19 | -2.78, 3.14 | 0.90 |
| Standard length | -1.07 | -2.27, 0.11 | 0.08 |

**Additional analyses and results: length of maternal exposure**

***Statistical analysis*.** Because males make their sperm at the beginning of the breeding season, we could remove them at set points in time and standardize exposure among males. However, because females took a variable amount of time under the predation treatment to become gravid (females lay clutches of ~50-150 eggs at a time), there was variation in the length of the predation treatment. We statistically test for the importance of this variation in two ways. First, we compare the full models (described in the original text) with and without maternal identity included, to understand if model fit was improved when the random effect of maternal identity was included. For survival models (linear mixed effects), we provide AIC values and statistical comparisons of the two models; for all other models using MCMCglmm, we provide DIC values (as with AIC values, lower scores indicate better model fit). There are no firm guidelines about relative differences in DIC scores, but generally, a difference of 5 is considered substantial (Spiegelhalter *et al.* 2002).

Second, we sought to understand if, among offspring with a predator-exposed mother, the number of days the mother was chased also altered offspring traits. Consequently, we reran the same statistical models as described above on the subset of offspring with a predator-exposed mother (all offspring with either just a predator-exposed mother or two predator-exposed parents). We retained all the fixed effects stated above (including paternal treatment), but replaced the binomial fixed effect of maternal treatment (exposed or control) with the fixed effect of the number of days the mother was exposed to the predator (continuous). We removed the random effect of maternal identity in these models, as it was redundant with the number of days chased. We report the statistical significance of this fixed effect of exposure length, as well as include AIC/DIC values for the model with and without this fixed effect.

***Results***

*Survival under predation risk*. In our full models, we found no detectable difference in model fit when maternal identity was included versus removed from the random effects (χ^2^=0.001, p=0.99; AIC with maternal identity: 422.35; without maternal identity 420.35). Among offspring with a predator-exposed mother, we found no evidence that the length of maternal exposure altered the likelihood that offspring were captured by the sculpin predator (Z_164_=0.33, p=0.74). We found no detectable difference in model fit when exposure length was included versus removed from the fixed effects (χ^2^=0.001, p=0.99; AIC with exposure length: 210.40; AIC without exposure length: 208.51).

*Stress-induced respiration*. In our full models, we found no improvement in model fit when maternal identity was included versus removed from the random effects (DIC with maternal identity: 7384.859; without maternal identity 7388.849). Among offspring with a predator-exposed mother, we found no detectable effect of length of maternal exposure on opercular beats (95% CI [-0.65, 0.84], p=0.80). We found no improvement in model fit when exposure length was included versus removed from the fixed effects (DIC with exposure length: 3699.714; DIC without exposure length: 3699.493).

*Behavioral assays.* In our full models, we found no improvement in model fit when maternal identity was included versus removed from the random effects for activity/exploration (DIC with maternal identity: 734.8107; without maternal identity 735.8208), freezing behavior (DIC with maternal identity: 910.7067; without maternal identity 910.83), length (DIC with maternal identity: 523.4239; without maternal identity 523.6977), or mass (DIC with maternal identity: -573.8096; without maternal identity -576.7375).

Among offspring with a predator-exposed mother, we found no detectable effect of the length of exposure on activity/exploration (95% CI [-0.03, 0.05], p=0.66), freezing behavior (95% CI [-0.04, 0.04], p=0.84), or mass (95% CI [-0.0003, 0.0003], p=0.84). We also found no improvement in model fit when exposure length was included versus removed from the fixed effects for activity/exploration (DIC with exposure length: 365.1608; DIC without exposure length: 363.8808), freezing behavior (DIC with exposure length: 464.7084; DIC without exposure length: 464.7604), or mass (DIC with exposure length: -268.829; DIC without exposure length: -269.3597). However, offspring of mothers who were chased longer were longer at 4.5 months compared to those whose mothers were chased for a shorter period of time (95% CI [0.03, 0.17], p=0.007) and model fits were improved when the length of exposure was included as a fixed effect (DIC with exposure length: 237.7696; DIC without exposure length: 243.0026).

*Scototaxis.* In our full models, we found no improvement in model fit when maternal identity was included versus removed from the random effects (DIC with maternal identity: 573.8661; without maternal identity 574.4545). Among offspring with a predator-exposed mother, we found no detectable effect of length of maternal exposure on scototaxis behavior (PC: 95% CI [-0.01, 0.05], p=0.18). We found no improvement in model fit when exposure length was included versus removed from the fixed effects (DIC with exposure length: 293.3457; DIC without exposure length: 293.504).
